# Family-Wide Photoproximity Profiling of Integrin Protein Social Networks in Cancer

**DOI:** 10.1101/2024.09.18.613588

**Authors:** Anthony J. Carlos, Dongbo Yang, Deborah M. Thomas, Shuyuan Huang, Keira I. Harter, Raymond E. Moellering

## Abstract

Integrin family transmembrane receptors mediate dynamic interactions between cells and their extracellular microenvironment. The heterogeneous interaction partners of integrins directly regulate cell adhesion, motility, proliferation, and intracellular signaling. Despite the recognized importance of protein-protein interactions and the formation of signaling hubs around integrins, the ability to detect and quantify these dynamic binding partners with high spatial and temporal resolution remains challenging. Here, we developed an integrin-family-directed quantitative photoproximity protein interaction (PhotoPPI) profiling method to detect and quantify native integrin-centered protein social networks on live cells and tissues without the need for genetic manipulation, antibodies, or non-physiologic cell culture conditions. We drafted quantitative maps of integrin-centered protein social networks, highlighting conserved and unique binding partners between different cell types and cellular microenvironments. Comparison of integrin social networks in cancer cell lines of diverse tissue of origin and disease state identified specific AND-gate binding partners involved cell migration, microenvironmental interactions and proliferation that serve as markers of tumor cell metastatic state. Finally, we identified unique combinations – or barcodes - of integrin-proximal proteins on the surface of pre- and post-metastatic triple negative breast cancer (TNBC) cells whose expression strongly correlate with both positive and negative disease progression and outcomes in TNBC patients. Taken together, these data provide the first family-wide high-resolution maps of native protein interactors on live cells and identify dynamic integrin-centered social networks as potential AND-gate markers of cell identity, microenvironmental context and disease state.

## Introduction

The cell surface proteome is a dynamic and heterogeneous landscape of proteins involved in myriad cellular processes. Among the many classes of proteins present on the cell surface, a subset is directly involved in communicating signals to and from neighboring cells and extracellular matrix components, thereby regulating intrinsic cellular responses to the extracellular environment. The integrin family of cell surface proteins is comprised of 18 distinct α-integrins and eight β-integrins, which form a network of 24 known functional heterodimers^1,2^. These transmembrane complexes mediate bidirectional, ‘inside-out’ and ‘outside-in’ signaling. ‘Outside-in’ signaling is typically induced by extracellular ligand binding, which facilitates cell spreading, retraction, migration, and proliferation, among other processes. Meanwhile, ‘inside-out’ signaling events are propagated by integrin cytoplasmic domains, which ultimately activate ligand binding and adhesiveness, as well as cellular migration^3–5^. Integrin heterodimer complexes are subclassified depending on their known ligand associations, including collagen receptors, laminin receptors, leukocyte-specific receptors, and ‘RGD’ receptors^6–8^. In cancer, integrins are implicated in several aspects of disease progression. The increased expression of integrins and activation of integrin signaling pathways contribute to proliferative phenotypes, local invasion and intravasation of metastasizing cancer cells, and survival of circulating tumor cells. Altered integrin activity is also implicated in the colonization, survival, and therapeutic resistance of metastatic tumor colonies^3, 9–11^. Despite the strong correlation between dysregulated integrin levels in cancer, relatively little is known about the protein-complex membership in integrin-centered ‘migratory hubs’ involved in bidirectional signaling with extracellular environments.

Previous studies have explored integrin interactors and signaling using a variety of canonical methods, including fluorescence microscopy, immunofluorescence, gene silencing, and affinity purification-mass spectrometry (AP-MS), amongst others^12^. These studies have identified tissue- and disease-specific integrin complex members^13^, as well as context-dependent interaction partners, including receptor tyrosine kinases like c-Src and EGFR^14, 15^. However, few chemical tools or methods exist for quantitative proteomic profiling of native integrins and integrin protein-protein interactions (PPIs) in live cells and tissues, mainly because quantitative detection of dynamic cell surface protein interactomes remains a significant technical challenge. Traditional AP-MS or immunoprecipitation approaches can overlook low affinity or transient protein interactions, fail to capture *in situ* protein interaction dynamics and raise the potential for false positive detection of interactions under non-physiologic lysis and pulldown conditions, among other challenges.

We hypothesized that many technical limitations could be overcome by the application of *in situ* ‘proximity profiling’ methods, which utilize locally activated reactive chemical tagging agents to label direct and proximal protein interactors.^16–20^ Enzymatic proximity profiling methods such as BioID, APEX, and PUP-IT rely on the expression of genetic fusions to a specific protein of interest, resulting in proximal binding partner labeling with reactive and retrievable chemical tags^21–24^. More recently, we described a light-activated proximity profiling approach, PhotoPPI, which enables mapping of intracellular and extracellular protein interactors through the local activation of reactive carbenes in response to a light trigger^16, 25^. This work, as well as subsequent approaches that use antibody conjugates of Ir- or small molecule-photocatalysts^26–28^ demonstrate the potential for minimally perturbative, rapid, and spatially restricted detection of protein interactors in or on live cells. We hypothesized that an ideal approach to profile native integrin ‘social networks’ would be to develop a similar technology without the need for genetic fusions, antibodies, or large-transition metal photocatalysts. Here, we describe the development of a small molecule ligand-directed photoproximity method to map native and heterogeneous integrins and their interactors on live cells and tissues. We demonstrate that this approach detects cell- and context-dependent integrin ‘social networks’ that serve as markers of cell state and are directly implicated in cancer pathology, such as patient survival in breast cancer.

## Results

### Design and Characterization of a Native Integrin-Directed Photoproximity Profiling Pipeline

To develop a photoproximity protein interaction (PhotoPPI) profiling method capable of capturing the interaction partners of native integrins on live cells, we first designed a fully synthetic chemical probe comprised of an ‘RGD’ peptide targeting motif, which is structurally similar to reported integrin-targeting agents^29–31^, fused to a synthetic lumichrome photocatalyst^32, 33^ (Fig. 1A). The RGD motif is a specific amino acid sequence naturally found on certain ECM proteins such as fibronectin, vitronectin, and laminin, which interacts with the RGD binding domains of more than half of all known integrins^34,35^. Therefore, an RGD-containing chemical probe should localize efficiently to diverse integrin receptor complexes on cell surfaces and present an opportunity for ‘family-wide’ profiling of the proximal ‘social network’ of neighboring surface receptors and ECM proteins. This information could not be captured through the use of single protein target-specific antibody conjugates or expression of genetic fusions, which have been the predominant approaches for *in situ* proximity profiling methods^16, 25–27^. To access a synthetically tractable and biologically compatible photocatalyst, we first designed a novel amine-reactive lumichrome photosensitizer, **1**, which should be broadly useful for conjugation to small molecules, peptides, and other biologics (Fig. 1A). We chose lumichrome due to its photostability and ability to activate a range of groups in response to light to generate reactive species^21^ for labeling of direct and proximal binding partners *in situ*. Here, we directly conjugated **1** to the RGD-targeting peptide core on solid-phase support, permitting direct cleavage and purification of the dual-function photoproximity probe, referred to hereafter as RGDL (Fig. 1A).

**Figure 1:**
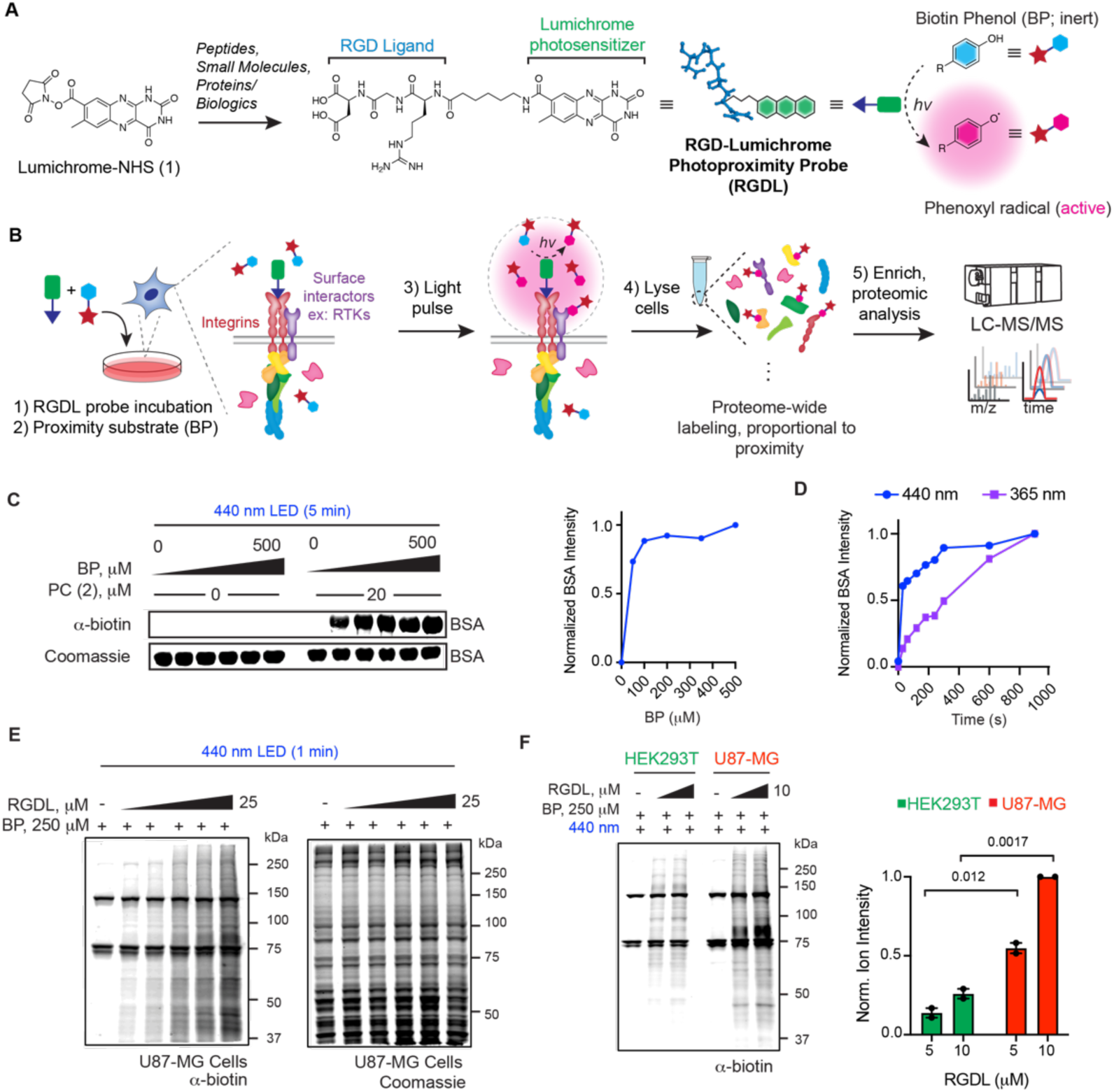
RGDL-dependent photoproximity profiling (RGDL-PhotoPPI) of native integrin complexes. **A**) Chemical structures of lumichrome-NHS (1) and RGDL photoproximity probe (*left*). Lumichrome functionalized chemical probes are used here to photosensitize an inert phenolic substrate into the reactive phenoxyl radical at RGDL-bound complexes *in situ* (*right*). **B**) Schematic of RGDL-PhotoPPI workflow on live cells. **C)** Detection of biotinylated BSA in response to RGDL workflow by immunoblot and Coomassie gel (*left*); quantification of BP substrate concentration-dependent biotinylation (*right*). **D**) Immunoblot quantification of time-dependent BSA photolabeling with 440 nm (blue) and 365 nm (purple) light activation. **E)** Anti-biotin immunoblot of U87-MG cells incubated with different RGDL ligand concentrations, followed by BP substrate treatment and blue light irradiation. **F**) Streptavidin-stained immunoblot (*left*) of RGDL photoprobed live U87-MG and HEK293T cell cultures. Quantification (*right*) of total lane intensity for each dose of RGDL tested. Data in C-F are representative samples from *n* = 2 biological replicates. P-values in F are from Student’s two-way t-test.

Our photoproximity profiling strategy aimed to localize RGDL molecules to native integrins on the cell surface where they can activate a spatially restricted, reactive substrate for proximal tagging of protein interactors and complexes (Fig. 1B). This strategy relies on the co-localization of a suitably reactive substrate for activation in the presence of light for spatial and temporal control of labeling events. To test this premise, we characterized the activation of a biotinamide phenolic substrate (referred to as BP) by the core photocatalyst, **2,** in a model solution-labeling assay using BSA. We observed strong BSA protein labeling at low micromolar concentrations of substrate (0-500 μm) and lumichrome photocatalyst (0-20 μM) in solutions exposed to either 365 or 440 nm light sources (Fig. 1C-D, Extended Data Fig. 1A-B). Photoactivation occurred rapidly (activation <1 min; Fig. 1D) and with very low or undetectable background in the absence of light or photocatalyst. Other substrates, including reported trifluoromethyldiazirines, could be activated by lumichrome but exhibited some light-dependent activation in the absence of photocatalyst and were thus ignored for the purposes here (Extended Data Fig. 1C-E). Finally, we performed experiments with RGDL, BP, and blue light activation directly on live U87-MG glioblastoma cells, demonstrating RGDL- and light-dependent biotinylation of native proteins under standard cell culture conditions (Fig. 1E). We next performed RGDL photolabeling on U87-MG and HEK293T cells, which representing relative high- and low-integrin expressing cell lines, respectively^36–39^, and observed significantly higher RGDL-dependent labeling in U87-MG cells (Fig. 1F). Collectively, these data supported the rapid and integrin-dependent specific photoproximity labeling with RGDL and blue light, providing conditions for further quantitative exploration by mass spectrometry-based proteomic profiling on cells and tissues.

### Quantitative RGDL-PhotoPPI Profiling of Unique and Conserved Integrin Complexes on Live Cancer Cells

Next, we sought to identify integrin-centered protein interactors on live cells using an RGDL-directed, quantitative photoproximity proteomic workflow. We first chose the U87-MG glioblastoma cells due to their relatively high expression of several RGD receptor integrins, such as ITGB1, ITGAV, ITGB3^37–39^, as well as previous use of RGD-derived probes and imaging agents^30, 40, 41^ with this cell line. Furthermore, despite being one of the most aggressive and common cancers that originate in the brain^42^, little is known about integrin-dependent protein complexes and signaling pathways in glioblastoma. To detect and quantify RGDL-labeled, and therefore integrin-proximal target proteins, we designed a PhotoPPI workflow using stable isotope labeling using amino acids in cell culture (SILAC) cell cultures. We pre-incubated matched U87-MG cultures with RGDL (‘heavy’ cultures) or vehicle alone (‘light’ cultures), washed off unbound ligand, and then added BP substrate followed by 440 nm light irradiation for 5 min (Fig. 2A). Under these conditions both cultures are exposed to substrate and light irradiation to control for any background, non-RGDL mediated protein detection in the workflow. After photolabeling, cells were lysed and processed for labeled protein enrichment and mass spectrometric analysis in a similar workflow previously developed for intracellular photoproximity profiling^25^.

**Figure 2:**
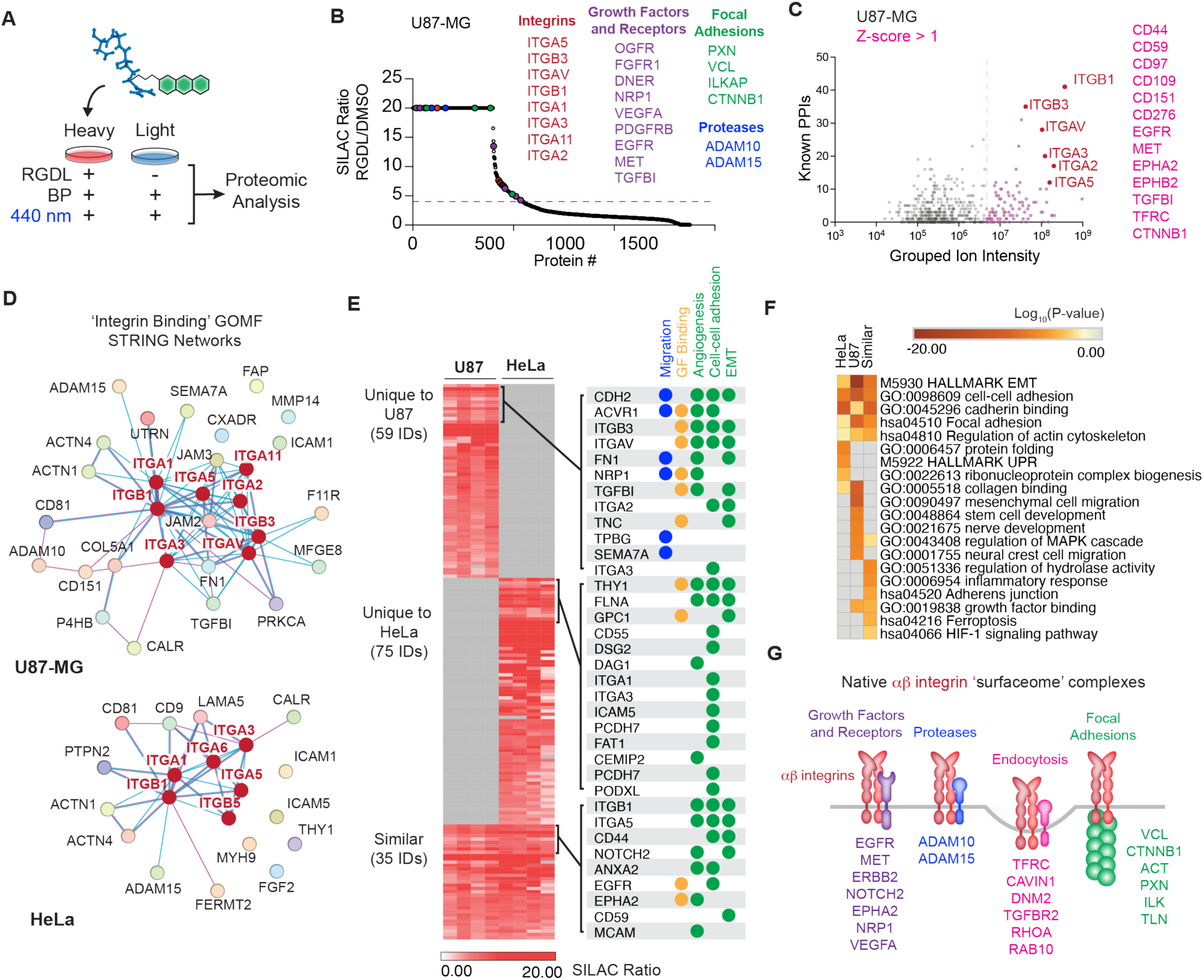
RGDL-PhotoPPI detects conserved and unique integrin interactomes on live glioblastoma and cervical cancer cells. **A)** Probe vs. no-probe RGDL-PhotoPPI target identification using SILAC-labeled cells. **B**) Waterfall plot of SILAC ratios (RGDL-treated/DMSO-treated) from U87-MG cells as described in (A). Raw enrichment of > 4-fold (i.e., RGDL probe-dependent labeling) in two or more biological replicates (from *n* = 4 total) was used as a cutoff for identifying proximal targets. Representative integrin (red), growth factor-related (purple), focal adhesion member (green), and surface protease (blue) proteins are highlighted. **C)** Yang plot depicting the relationship between the level of enrichment (median ion intensity) and the number of STRING database-annotated protein-protein interactions (PPIs) for each protein enriched from analysis in (B). Highly enriched proteins with the most abundant connectivity appear in the top right of the plot. Integrins (red) and proteins that are one standard deviation (SD) above the mean log_10_(‘Heavy’ grouped ion intensity; pink) are highlighted. **D**) U87-MG (*top*) and HeLa (*bottom*) connectivity plots of enriched proteins from the ‘integrin binding’ gene ontology molecular function (GOMF) group. **E**) Heatmap depiction of unique and common integrin interaction partners identified in (B-C) from U87-MG and HeLa (*left*), and memberships within select pathways (*right*). A subset of enriched proteins is shown on the right. **F**) Heatmap of unique and conserved enriched GO, KEGG, and HALLMARK pathway terms from Metascape analysis of the data in (E). **G**) Graphic depiction of some conserved αβ-integrin social network members identified by RGDL-PhotoPPI profiling in U87-MG and HeLa cancer cell lines.

Top enriched targets, both in terms of SILAC ratio (i.e., significant labeling in an RGDL-dependent manner) and median ion abundance were native integrins, including several RGD-type receptor integrins such as ITGB1, ITGAV, ITGB3, and ITGA5. These top hits form known RGD-specific αvβ3 and α5β1 heterodimeric receptors (Fig. 2B-C)^31^. In addition, we were able to identify several laminin and collagen receptor integrins (ITGA1, ITGA2, ITGA11, ITGA3), presumably due to either direct probe interaction or close proximity to cell-surface signaling hubs containing RGD-binding integrins (Figure 2B-C). Beyond labeling core integrins, RGDL labeling detected numerous proteins that are known integrin-associated interactors, including several growth factor receptors and receptor tyrosine kinases such as EGFR^43^, MET^44^, FGFR1^45^, ADAM-family cell surface proteases^46^, as well as members of integrin-anchored focal adhesion complexes (e.g., VCL^47^, PXL^48^ and ACTN^49^; Fig. 2B-C). To assess the relative strength or efficiency of integrin interactions beyond probe-dependent enrichment alone (i.e., SILAC ratio between probe and no-probe conditions), we plotted the median ion intensity of enriched targets across biological replicates and performed an interaction-based STRING analysis of all enriched targets within the enriched target protein set from U87-MG cells (Fig. 2C-D). In the latter analysis, we plotted the total known interactions for a given target protein with any other enriched protein in the dataset based on STRING databases from the literature (hereafter referred to as a Yang Plot). We observed that several core integrin members and known and novel associated surface receptors like EGFR, CD44, TGFB1, EPHA2, and MET were enriched in the upper right quadrant of the Yang Plot (Fig. 2C). Combined, we posit that proteins demonstrating RGDL-dependent enrichment and significantly higher abundance and connectivity may represent core interactors with integrin complexes in these contexts.

With this enrichment workflow in place, we next profiled the integrin social network in HeLa cells, a well-studied cervical cancer cell line with relatively lower expression of several integrin receptors such as ITGB1 and ITGB3^37, 38^. As in U87-MG cells, RGDL-PhotoPPI profiling identified several integrins (ITGB1, ITGA5, ITGA3, IGTB5 and others), surface receptors, proteases, and adhesion complex members (Fig 2D-E; Extended Data Fig. 2A) as enriched and abundant complex members. Comparative analysis of enriched target proteins between U87-MG and HeLa cells revealed similarities and differences. Integrin ‘social networks’ on both cells contained proteins involved in cell-cell interactions, cadherin binding, and epithelial-mesenchymal-transition (EMT)-associated pathways (Fig. 2E-F). The quantitative nature of RGDL-PhotoPPI enabled the identification of distinct pathways and proteins across cell types. This included enrichment of neural crest and stem cell-related proteins in the glia-derived U87-MG cells, as well as increased enrichment of proteins involved in cell migration and angiogenesis. By contrast, less invasive HeLa cells contained more proximal proteins involved in endocytosis, protein folding, and specific cadherin family members (Fig. 2E-F). These comparative profiling analyses are congruent with previous work indicating that highly invasive U87-MG glioblastoma cells are dependent on integrin-mediated interactions, thus enabling development of diagnostic and therapeutic agents targeting integrins in this tumor type^50,51^. Finally, these data collectively suggest that RGDL-directed profiling on live cells can capture integrin interactors across various points in their lifecycle, including endocytosis, internalization, and perhaps receptor recycling (Fig. 2G) and that these interaction networks have unique membership associated with cell type and cell state.

### Cellular Environment Significantly Alters Integrin Social Network Membership

Beyond comparing integrin social networks between disparate cell types, we hypothesized that the flexibility of RGDL-dependent photoproximity profiling would permit interrogation of altered integrin interaction networks in response to changes in the extracellular environment. To test this hypothesis, we probed the dynamic changes to integrin interaction wiring between adherent and detached U87-MG and HeLa cells, mirroring the populations of cells that are in contact with extracellular matrix versus those that have detached (i.e., undergoing metastasis or anoikis). While many studies have explored signaling changes or specific interactions involved in cell detachment and dissemination^52^ little is known about the changes to integrin-centered interaction networks under these and many other context-dependent conditions.

Using matched adherent and detached cell cultures, we performed RGDL-dependent photoproximity labeling, enrichment, mass spectrometry analysis, and quantitative comparison of interaction networks between each population (Fig. 3A). Principal component analysis confirmed clustering among cell type- and condition-matched replicates, which showed that altered extracellular environment had a more pronounced effect on the overall integrin social network than cell type (Fig. 3B). Integrins remained amongst the most abundant enriched target proteins in each cell line and under both adherent and detached conditions (Fig. 3C-E, Extended Data Fig. 2A-B). In addition to some integrins, there were interacting proteins that were equally enriched between adherent/detached cultures, such as ICAM5, but in general the core integrin-centered network and overall connectivity within these networks were reduced in detached U87-MG and HeLa cells (Fig. 3D-E). We performed gene ontology (GO) analyses on detached and adherent cultures in both cell lines, which highlighted increased enrichment of ‘integrin binding’ and ‘receptor protein tyrosine kinase activity’ GO molecular function terms, among others, in adherent cells (Fig. 3F, Extended Data Fig. 2C). Similar analysis of enriched GO cellular component terms were consistent with increased internalization of integrins^53^, and a strong reduction in the enrichment of GO terms associated with cell-cell interaction, tight junctions, and extracellular matrix interactions in detached cells (Fig. 3F, Extended Data Fig. 2C). These data collectively demonstrate the ability of the RGDL platform to label and detect conserved and unique integrins, integrin PPIs, and integrin-dependent pathways across cancer cell lines of varied origin and microenvironmental context.

**Figure 3:**
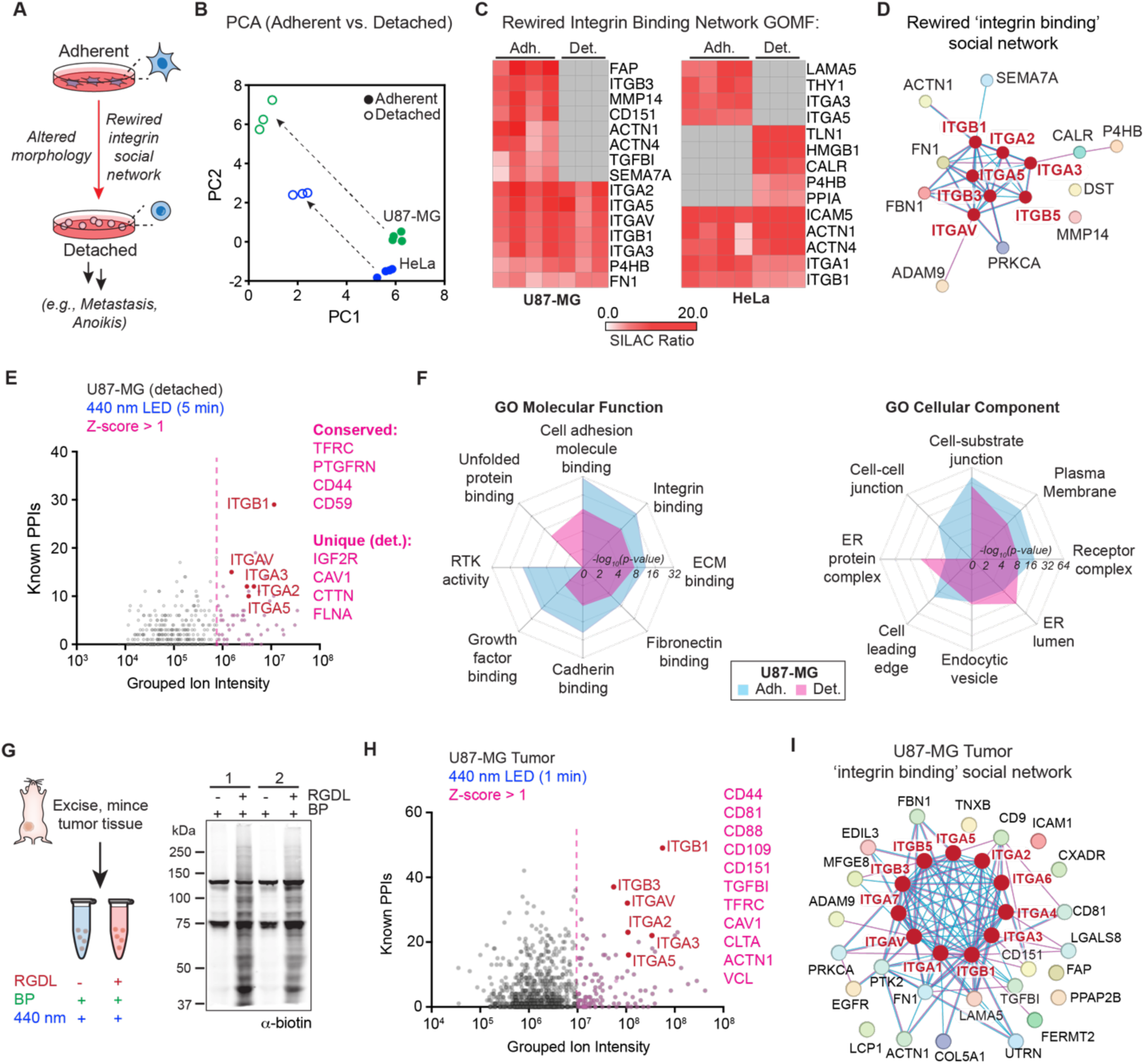
RGDL-PhotoPPI mapping in diverse cellular contexts. **A**) Schematic of attached and detached conditions used for RGDL-PhotoPPI profiling. **B**) Principal component analysis (PCA) of integrin interacting members in adherent and detached cultures of HeLa and U87-MG cells. **C**) Heatmaps showing a subset of integrin-binding associated protein ratios that are consistent and unique to adherent and detached cultures. **D-E**) GOMF ‘integrin binding’ STRING network (D) and Yang Plot (E) of RGDL-enriched proteins from detached U87-MG cells. **F)** Radar plots comparing gene set enrichment P-values of attached and detached U87-MG cells. GO Molecular Function terms (*left*) and GO Cellular Component terms (*right*). Data showing in B-F are from *n* = 3 biological replicates for detached U87-MG and HeLa cell cultures. **G**) Schematic of RGDL-PhotoPPI tissue profiling of mouse tumor xenografts (*left*) and anti-biotin immunoblot analysis of MDA-MB-231 xenografts treated with RGDL probe and processed as indicated from *n* = 2 biological replicates (*right*). **H-I**) Yang Plot (H) of RGDL-PhotoPPI-enriched proteins from U87-MG tumor xenografts and STRING network (I) of proteins in the ‘integrin binding’ gene ontology molecular function (GOMF) category.

### RGDL-PhotoPPI Profiling on Native Tissue

We next sought to test whether the RGDL-PhotoPPI workflow is compatible with clinically relevant cell lines and primary tissues, starting with intact tumor tissues. Triple-negative breast cancer (MDA-MB-231) and glioblastoma (U87-MG) tumor xenografts were established in mice and excised upon reaching ∼300-500 mm^3^. Tumors were processed into smaller sections and processed for RGDL-PhotoPPI as described above for cell lines (Fig. 3G, Extended Data Fig. 3A). As with cell line studies, we confirmed protein labeling only in the presence of RGDL probe on MDA-MB-231 tumors, as determined by Western blot, matching previous results with cell lines (Fig. 3G). We then performed label-free quantification (LFQ) LC-MS/MS measurements following streptavidin-biotin enrichment to identify the integrin complex targets on U87-MG and MDA-MB-231 tumor tissues treated with 1 or 5 minutes of light, respectively (Fig. 3H, Extended Data Fig. 3 B-C). Notably, the U87-MG-derived xenograft subjected to minimal irradiation resulted in robust enrichment of the primary integrin network (i.e., ITGB1, ITGB3) and established peripheral social network members (i.e., CD44, TGFBI, CAV1, CLTA, EGFR). Generally, experiments with both xenograft types resulted in a more robust enrichment of target integrins and integrin-binding PPIs, as compared with cell culture experiments. Furthermore, the tumor-derived integrin binding networks are highly interconnected and seemingly more complex (Fig. 3I, Extended Data Fig. 3C); this may be the result of greatly increased cell-cell interactions within the tissue matrix compared to two-dimensional cell cultures. Overall, these data confirm that RGDL profiling can detect integrin-centered protein interactions on small (sub-mg) samples of primary tumor tissues and a facile workflow that could be applied to primary human tissues in future studies.

### Integrin Social Networks Correlate with Disease State and Outcomes

We next asked whether RGDL photoproximity profiling could identify interactions associated with disease states, such as cancer metastasis. The identification and characterization of surfaceome markers of metastasis, along with their use as prognostic markers or drug targets, could offer options for tailored therapy to improve patient outcomes. To determine how integrin social networks might change when cancer cells migrate from the primary site to a metastatic niche, we performed quantitative RGDL profiling on two closely related triple-negative breast cancer (TNBC) cell lines, MDA-MB-231 (231) and BM1.

The MDA-MB-231 cell line was developed from a pleural effusion collected from a TNBC patient. The metastatic BM1 cell line was developed by establishing MDA-MB-231 xenografts in immunodeficient mice, waiting for spontaneous metastasis and subsequent cell line generation from isolated bone metastases^54^. RGDL profiling was performed with SILAC-labeled pairs of 231 and BM1 cell cultures as described for adherent cultures, followed by quantitative proteomics (Fig. 4A). While whole proteome profiling demonstrated relatively uniform integrin abundance between the two cell types (Extended Data Fig. 5A-B), RGDL interaction profiling identified 175 and 680 integrin interactors that were significantly increased in 231 and BM1 cells, respectively (Fig. 4B, Extended Data Fig. 4A-B). STRING bioinformatic analysis of enriched target proteins highlighted that core integrins (e.g., ITGB3, ITGA2, ITGAV, and ITGB1) and previously identified integrin interacting proteins (e.g., CD44, EGFR and MET) were shared in both cell lines. Likewise, most of the general pathways associated with enriched interactors were shared, including those related to migration, adhesion, proliferation, regulation of apoptosis, and angiogenesis (Extended Data Fig. 4C). Within these networks there were many known and novel integrin-interacting proteins enriched. For example, ITGB3 and EGFR show similar probe enrichment values (i.e., SILAC ratio) and integrated ion intensities in both 231 and BM1 cells. To try and capture integrin proximity and labeling efficiency we calculated the product of these two as an integrated ‘proximity score,’ which likewise showed similar presence of ITGB3 and EGFR in both cell lines (Extended Data Fig. 4D). Other known interactors were differentially enriched in the two cell lines. For example, VCAM1, a known integrin interactor^55^ and potential drug target in metastatic breast cancer^56^ was highly enriched in 231 but not in post-metastatic BM1 cells (Fig. 4C), By contrast, ICAM2, a ligand of ITGB2 was only enriched in BM1 cells, which is not surprising given its implication as a promoter of brain metastasis and tumor cell stemness in TNBC^57^. SLC3A2, a known interactor of ITGB1^58^, is overexpressed in many cancer types, including breast cancers, and has been proposed as a drug target^59, 60^. We observed that SLC3A2 isoform 1 was uniquely enriched in 231 cells, while isoform 2 was found only in BM1 cells (Fig. 4C), highlighting isoform-specific complex tracking of cancer-associated targets with RGDL profiling.

**Figure 4:**
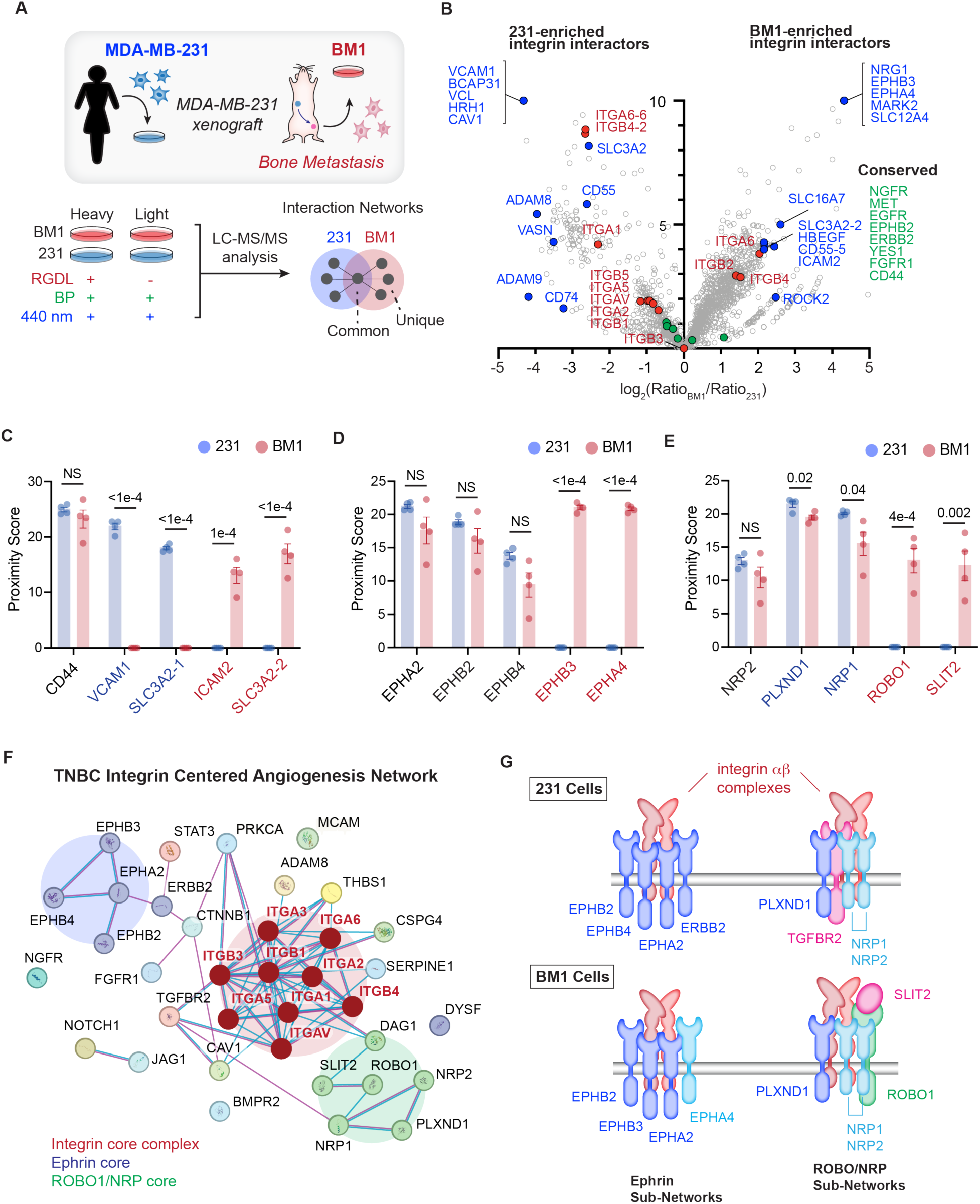
Unique integrin interaction networks typify TNBC cell line metastatic stage. **A**) Schematic depiction of MDA-MB-231 and BM1 cell line origin (*top*) and SILAC-based RGDL-PhotoPPI profiling of surface interactors therein. **B**) Volcano plot comparing relative abundance (via RGDL-PhotoPPI SILAC ratio) of integrins and interaction partners on live MDA-MB-231 and BM1 cells; *n* = 4 biological replicates. **C-E**) Quantitative Proximity Scores in 231 and BM1 cells for previously known integrin interacting proteins (C) and novel integrin interactors in the Ephrin (D) and NRP/ROBO (E) receptor family ratios. **F)** STRING network depiction of angiogenesis-related integrin interactors detected in 231 and BM1 cells. **G**) Schematic depiction of differential Ephrin- and NRP/ROBO sub-network interactions with αβ-integrin-complexes in MDA-MB-231 and BM1 cells. Data in (B-E) represent mean ± S.E.M. from *n* = 4 biological replicates. Statistical comparisons are two-tailed unpaired *t-*tests.

Beyond known interactors, RGDL profiles highlighted many novel integrin interacting proteins that were differentially present in BM1 and 231 cells. For example, proteins related to angiogenesis were among the most significantly enriched integrin interactors in both cell lines (Fig. 4D-F). The angiogenic network was centered on an integrin core, with sub-networks including ephrin receptors and ROBO1/SLIT signaling complexes. For example, while both cell lines show enrichment of neuropilins (NRP1 or NRP2), post-metastatic BM1 cells show unique proximity between both ROBO1 receptor and its ligand SLIT2 (Fig. 4E-F), which is significant, as the ROBO1-SLIT2 axis has been shown to promote metastasis in pancreatic cancer^61^. While ROBO-SLIT complexes are annotated as potential integrin interactors at the pathway level, no direct interactions at the PPI level have been previously documented^62^. Additionally, BM1 and 231 cells have both shared and unique ephrin receptors, which are receptor tyrosine kinases strongly implicated in cancer (Fig. 4D, F)^63^. For example, EPHA2 and EPHB2 were proximal to integrins at roughly similar levels in 231 and BM1 cells, but other ephrins showed differential proximity across cell types. EPHB4 integrin proximity was increased in 231 cells, whereas EPHA4 and EPHB3 integrin proximity was uniquely detected in post-metastatic BM1 cells (Fig. 4D, F), matching their previously identified roles in promoting cancer progression^64–67^. These data present the potential for quantitative and comparative RGDL-PhotoPPI profiling to identify ‘AND-gate’ complex pairs (i.e., the proximity of integrins and identified surface markers on live cells, Fig. 4G) that could serve as novel players in breast cancer metastasis and post-metastatic colonization, as well as potential markers for diagnostic or therapeutic interventions.

To evaluate whether the unique integrin social network members might serve as prognostic markers, we assembled a 20-protein signature, or ‘interaction barcode,’ for BM1 (metastatic) and MDA-MB-231 cells (primary) by selecting the most abundant and unique surface proteins within the enriched RGDL-labeled integrin social network in each cell line (Extended Data Fig. 4E). Since broad-scale clinical databases with matched protein interaction partner information do not exist, we hypothesized that the mRNA expression of these targets within annotated breast cancer patient databases could serve as a pragmatic - albeit imperfect - proxy. Therefore, we performed retrospective survival analyses of breast cancer patients by comparing relative mRNA expression levels of each interaction partner signature (Fig. 5A) within published databases of TNBC and breast cancer patient populations^68, 69^ (Fig. 5A-B). Surprisingly, we found that TNBC patients with higher mRNA expression levels of the primary tumor barcode (e.g., 231 cell-derived integrin partners) had a significant survival advantage as compared to patients with lower expression levels (Fig. 5A; Hazard Ratio, HR = 0.5, *p* = 0.008). Expression of the metastatic integrin interactor barcode (e.g., those unique to BM1 cells) was significantly correlated with increased mortality in TNBC patients (HR = 1.92, *p* = 0.01, Fig. 5A). These trends held true even amongst a larger cohort of 1,089 patients with diverse forms of breast cancer (HR = 0.51 and HR = 1.63 for the primary and metastatic barcodes, respectively; Fig. 5B), thus unveiling significant correlations between integrin social networks of metastatic cells – identified through unbiased RGDL-PhotoPPI profiling – and patient outcomes. Collectively, these data may offer new avenues for targeted therapy and prognostic marker development in aggressive breast cancer types.

**Figure 5:**
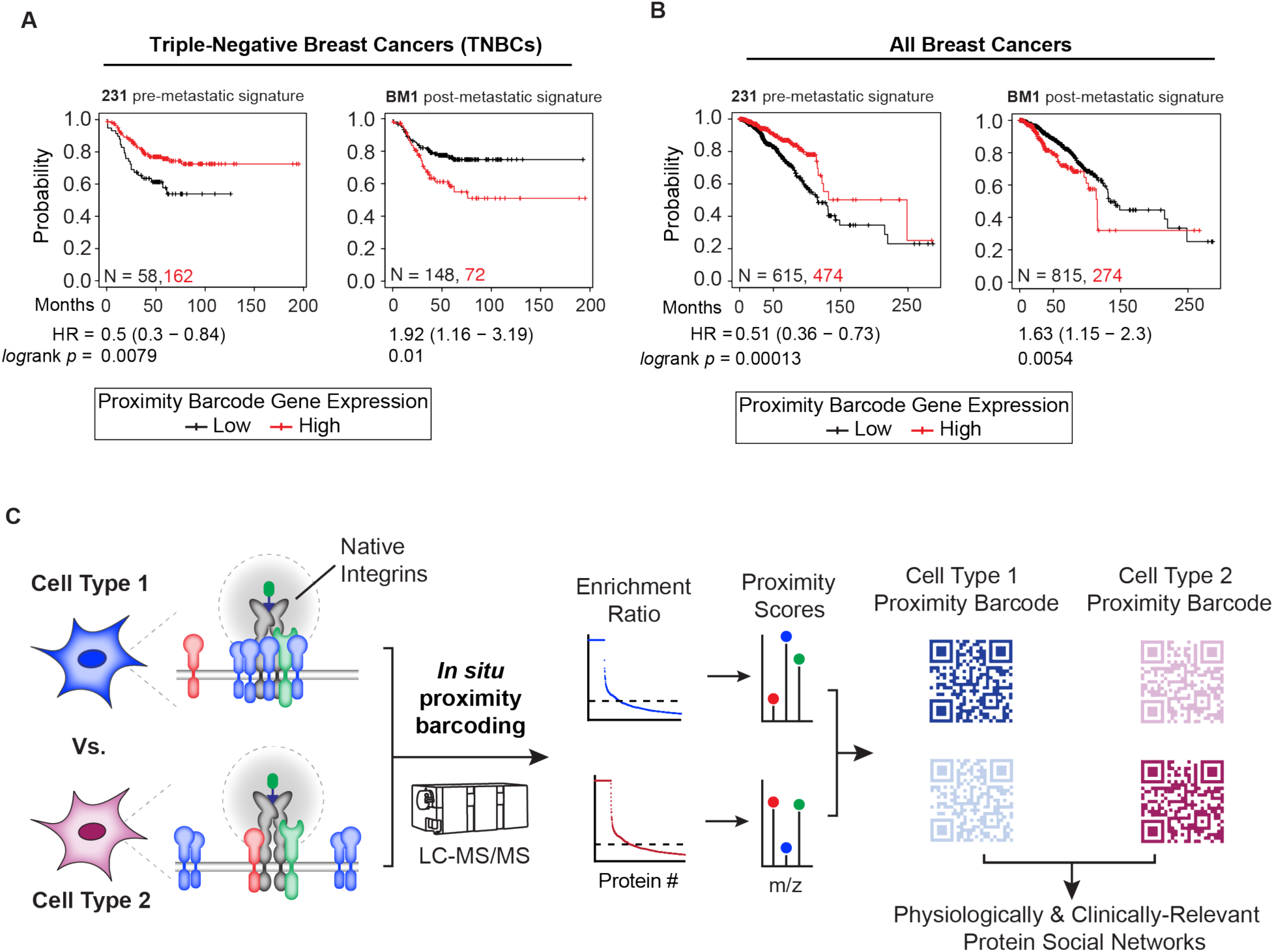
Expression of primary- or metastatic tumor-specific Integrin interactors correlates with patient outcomes in human breast cancers. **A**) Kaplan-Meier (KM) analysis of triple-negative breast cancer (TNBC) patient survival based on relative mRNA expression levels of unique 20-gene signatures derived from MDA-MB-231 cells (*left*) or BM1 cells (*right*). Survival data of patients with higher aggregate barcode expression are represented in red, and those grouped by lower expression are in black. **B**) KM analysis of patients from all breast cancers in the KM database. Hazard ratios (HR) with 95% confidence intervals and logrank P-values are indicated below each survival plot in A and B. **C**) RGDL-PhotoPPI for the *in situ* ligand directed proximity labeling of native integrins to identify disease relevant protein interactome barcodes between cell types.

## Discussion

In this study, we aimed to develop a photoproximity profiling method capable of interrogating integrin-centered protein interactors, or ‘social networks,’ directly on live cells. To do so, we developed an integrin sub-family targeting RGDL probe that can localize to and proximally activate a reactive phenoxyl radical tagging species in response to light. We demonstrated several advantages to this approach. First, by bypassing the use of genetic fusion proteins and/or antibody-based recognition, RGDL-PhotoPPI can probe heterogeneous, native integrin complexes that occur on live cells and tissues. *In situ* labeled protein retrieval, detection, and quantification by MS-based proteomics enables unbiased discovery of interacting target proteins without *a priori* information about their identities. Moreover, this approach allows for qualitative and quantitative comparison of integrin social networks between diverse tumor cell types, microenvironmental contexts, and metastatic status. Studies of surface complex dynamics and the impact of microenvironmental conditions, in particular, can be sensitive to external stimuli, and therefore the use of rapid light activation as a proximity trigger is minimally perturbative relative to other proximity profiling approaches. The speed of RGDL-PhotoPPI photoactivation is also noteworthy, demonstrated by robust proteome coverage and high enrichment over background following irradiation times as low as one minute. Future studies could leverage this temporal resolution to detect dynamic integrin interactors across their lifecycle and in different sub-cellular locales following defined perturbations.

While it is commonly understood that integrin signaling plays a role in basal cellular function, our results here revealed significant differences in integrin interactomes across different cancer cell lines and contexts. RGDL-PhotoPPI detected numerous previously validated integrin interactors, including cell surface growth factor receptors like EGFR, ERBB2 and MET, as well as known and novel proteins involved in ECM remodeling, focal adhesion complexes and extracellular proteases (Fig. 2). Within these larger protein groups, we identified considerable differences between cell types. For instance, integrin networks were enriched for endosomal and focal adhesion complex interactors in epithelial cervical cancer (HeLa) cells relative to the invasive glioblastoma (U87-MG) cells, which showed higher enrichment for interaction partners implicated in proliferative signaling, migration, and ECM remodeling. Intriguingly, we detected significant rewiring of integrin-centered social networks when comparing adherent versus detached cell populations from HeLa and U87-MG tumor cells. This result is not entirely surprising, given the role of integrins in ECM interactions and migration, however it raises the possibility of mapping and perhaps targeting distinct extracellular complexes that differ based on tumor cell microenvironments. Coupled with the ability to directly profile fresh tumor samples, we posit that RGDL-PhotoPPI could be applied in the future to map cell surface migratory hubs more broadly across normal and diseased tissue samples.

The sensitive and quantitative detection of integrin social networks is perhaps most useful to identify ‘AND-gate’ protein complexes that differentiate cells of different disease potential, such as the pre- and post-metastatic TNBC cell lines. While our comparison of 231 and BM1 cells confirmed that many integrin targets and interactors are similar, we detected many novel interactors previously associated with cancer progression and metastasis that were uniquely present in one cell line. The detected Ephrin and NRP/ROBO sub-complexes that are present in either pre- or post-metastatic TNBC cells provide potential hypotheses connecting cell migratory and signaling complexes co-localizing and perhaps forming synergistic ‘signalosomes.’ Further investigation is needed to elucidate the roles of these newly identified interactions in cancer progression and metastasis.

Finally, we used photoproximity profiling for the first time to identify metastasis-association interaction ‘barcode’ protein sets (Figure 5C). Intriguingly, the application of these social networks to clinical databases uncovered significant survival advantages for patients with higher expression of the pre-metastatic interactors (derived from MDA-MB-231 cells) and significantly reduced survival among patients with higher expression of post-metastatic interactors (derived from BM1 cells). These results raise the intriguing possibility that combinations of receptors, even when anchored around seemingly ubiquitous surface proteins like integrins, could serve as markers of and potential targets for tumor metastasis or other disease states. For example, the identification of select integrin-proximity of proteins like VCAM1, which has been targeted in breast cancer, in 231 vs. BM1 cells could suggest differential therapeutic efficacy based on breast cancer stage and support diagnostic profiling by RGDL-PhotoPPI as a method to confirm therapeutic potential for certain patients or cancer sub-types. Coupled with the ability to directly profile interactors on fresh tissues, this raises the possibility of RGDL-PhotoPPI profiling of patient samples for further discovery of select interaction markers with potential diagnostic utility. A caveat with and limitation of these data is that clinical databases of integrin-centered or other surface receptor-centered interactions are not available for review, requiring mRNA levels to be queried here as a proxy. Taken together, we posit that further development of proximity profiling methods like RGDL-PhotoPPI will provide opportunities to draft high-quality, spatially-resolved maps of integrin and other protein interaction social networks directly in physiologically and disease-relevant contexts.

## Methods

### General synthetic methods

Reagents purchased from commercial suppliers were analytical grade and used without further purification. All reactions were carried out in oven-dried flasks, using oven-dried magnetic stir rods and anhydrous solvents (Acros) unless otherwise specified. Nuclear magnetic resonance spectra were acquired using either a Bruker AVANCE II+ 500; 11.7 Tesla NMR or Bruker DRX 400; 9.3 Tesla NMR instrument. Accurate mass measurements and final probe purification were obtained and performed using an Agilent 1290 Infinity II with an Agilent 5 Prep C18 50 x 21.2mm column.

### Cell culture

The U87-MG cell line was obtained from the University of Chicago Cellular Screening Center. MDA-MB-231 and BM1 cells were gifted by Marsha Rosner (University of Chicago). The MDA-MB-231 cell line was originally obtained from ATCC and the BM1 cell line was generated by Kang and colleagues^54^ and obtained from Joan Massagué (Memorial Sloan-Kettering Cancer Center).

U87-MG and HeLa cells were propagated in RPMI (Corning) supplemented with 10% fetal bovine serum (FBS, Corning) and 1% penicillin/streptomycin (Gibco). MDA-MB-231 and BM1 lines were propagated in DMEM (Invitrogen). All cell lines were grown at 37 °C in a 5% CO_2_ humidified incubator. All cell lines tested negative for mycoplasma using the Lonza MycoAlert PLUS Detection Kit.

### SDS-PAGE and Immunoblot

Cells were lysed by on-well lysis using ice-cold 1 x RIPA (Millipore) and tip sonicated (Fisher Scientific FB-505) over ice. Insoluble debris was cleared by centrifugation, and the supernatant was diluted into 4X Laemmli buffer containing 50 mM dithiothreitol (DTT) as a reducing agent. Samples were prepared for SDS-PAGE by heating to 95 °C for 5 min, cooled to room temperature, resolved on 10% SDS-PAGE gel, and transferred to nitrocellulose membranes by standard western blotting methods. Membranes were blocked in 2% BSA in TBS containing 0.1% tween-20 (TBST) and probed with streptavidin-800 (Licor Odyssey CLx) to visualize biotin labeling. Blot intensities were quantified in Image J and normalized in Microsoft Excel.

### *In vitro* photosensitization experiments

A solution of BSA (10 μM) in PBS was prepared for all *in vitro* photolabeling assays. The optimal concentration of photocatalyst for *in vitro* photolabeling experiments was determined by performing a dose gradient of photocatalyst **2** (0-20 μM) on solutions of BSA with 100 μM BP with irradiation at 440 nm LED for 5 min. For substrate dosing assays, samples were treated with vehicle or lumichrome photocatalyst **2** (20 μM) from 5 mM stock solution. BP (0, 50,100, 200, 350, 500 μM) was then added and samples were then irradiated with a Kessil PR160L (LED, 440 nm) or Spectroline XL-1500 (365 nm) for 5 min on ice. For photokinetic experiments, BSA solutions with 100 μM BP and 20 μM photocatalyst **2** were irradiated with either wavelength for 0, 30, 60, 120, 180, 240, 300, 600, or 900 s, with samples collected at each time point. Samples were added to 4x loading buffer, vortexed, and boiled before immunoblot and Coomassie stain.

### RGDL probe dosing on live cells for immunoblot analysis

Wildtype U87-MG cells were seeded in 12-well plates at 250,000 cells per well in 1.5 mL of RPMI media (Corning) supplemented with 10% fetal bovine serum (Corning) and 1% penicillin/streptomycin (Gibco) 24 hours before the experiment. Cells were treated with 0, .5, 1, 5, 10 and 25 μM of RGDL in 500 μL of serum-free, phenol-red-free RPMI media (Gibco) for 1 h at 37 °C. After incubation, media was replaced with 500 μL of PBS with RGDL (0-25 μM) and BP (250 μM). The cells were irradiated under a Kessil PR160L (LED, 440 nm) for 1 min on ice. The cells were then washed in 1 mL of PBS twice and lysed in ice-cold 1x RIPA buffer supplemented with protease inhibitor (Roche). Cells were lysed while shaking on ice for 20 min before transfer and tip sonication (3 x 1 s pulses at 30% amplitude). The lysates were collected and processed for anti-biotin immunoblot analysis.

For comparative experiments between high and low integrin expressing cell lines, U87-MG and HEK293T cells were each seeded in 12-well plates at 250,000 cells in 1.5 mL of RPMI 48 h and 24 h before experimentation for U87-MG and HEK293T, respectively. Cells were treated with 5 and 10 μM of RGDL in 500 μL of serum-free, phenol-red-free RPMI media for 1 h at 37 °C. Cells were then irradiated and processed as indicated above. Lysates were then normalized to 1 mg/ml using BCA Protein Assay Kit (Pierce) and processed for anti-biotin immunoblot analysis. Total lane intensities were quantified in Image J and normalized to the most intense lane. Statistical comparison of intensities between each dose of probe between cells lines was performed in Graph Pad Prism using multiple unpaired t-test.

### SILAC cell culture methods and proteomic sample preparation

SILAC labeling was performed by growing cells for at least five passages in lysine- and arginine-free SILAC medium (RPMI or DMEM, Invitrogen) supplemented with 10% dialyzed fetal calf serum (Gemini) and 1% Pen/Strep. “Light” and “heavy” media were supplemented with natural lysine and arginine (0.1 mg/mL) for “light”, and ^13^C-, ^15^N- labeled lysine and arginine (0.1 mg/mL) for “heavy”, respectively.

### Sample preparation and streptavidin enrichment

Quantitative proximity labeling studies with SILAC quantitative proteomics were performed with “heavy” and “light” labeled U87-MG, HeLa, MDA-MB-231, and BM1 cell lines. SILAC-labeled cells, grown to 80-90% confluency in 10 cm cell-culture treated plates each, were incubated with DMSO alone (light cells) or RGDL probe (50 μM, heavy cells) for 1 h in serum-free SILAC RPMI. After incubation, excess probe was removed by aspiration and supplanted with BP (250 μM) in DPBS. Cells were then irradiated using a Kessil PR160L (LED, 440 nm) for 5 min, rinsed, scraped, and washed with cold PBS (1 ml x 4). The cells were pelleted and then lysed in RIPA lysis buffer (50 mM Tris, 150 mM NaCl, 1% Triton X-100, 0.5% deoxycholate, pH 7.4) supplemented with EDTA-free complete protease inhibitor (Roche) and 1 mM DTT, at 4 °C. After sonication, insoluble debris was cleared by centrifugation (17,000 *g*, 15 min). BCA assay was performed to normalize Heavy and Light protein concentrations to ∼1 mg/ml. Streptavidin agarose beads (50 μL slurry, Pierce) were washed twice with RIPA buffer, and each cell lysate was separately incubated with the beads with rotation overnight at 4 °C. The beads were subsequently washed five times with 0.5 mL of RIPA lysis buffer containing 1 mM DTT, combined together, then washed once with 1 mL of 1 M KCl, four times with 0.5 mL PBS, and two times with 2 M Urea in 25 mM ammonium bicarbonate. 500 μL of 6 M Urea in 50 mM ammonium bicarbonate was then added to the beads, and samples were reduced on resin by TCEP (10 mM final) with orbital shaking for 20 minutes at 65 °C. Samples were then alkylated by adding iodoacetamide (20 mM final), covered from the light and with orbital shaking, for 40 minutes at 37 °C. The streptavidin agarose beads were collected, washed once with 2 M Urea in 25 mM ammonium bicarbonate, and the buffer exchanged to 2 M Urea in 25 mM ammonium bicarbonate supplemented with 1 mM CaCl_2_. Enriched proteins were digested on-bead by the incubation of 2 μg sequencing grade trypsin overnight at 37 °C. Following trypsinization, supernatant was collected, acidified with HPLC grade formic acid (2% final, pH 2-3), and peptides were then desalted on ZipTip C18 tips (100 μL, Millipore), dried under vacuum, resuspended with LC-MS grade water (Sigma Aldrich), and then lyophilized. Lyophilized peptides were dissolved in LC-MS/MS Buffer (H_2_O with 0.1% formic acid, LC-MS grade, Sigma Aldrich) for proteomic analysis.

### Cell detachment studies

U87-MG and HeLa Cells were grown on 10 cm dishes to ∼80-90% confluency, rinsed with PBS and incubated with Cell Stripper non-enzymatic dissociation buffer (Corning, 5 mL, 37 °C) for 10 min and transferred to Falcon tubes. Cells were then pelleted (300 g, rt) and washed once with serum-free RPMI (-phenol red). Cells were then resuspended in RGDL-probe containing media (50 μM) or DMSO for 1 h in 4 mL of SILAC media, followed by replacement with DPBS containing BP substrate (250 μM). Cells in suspension were then irradiated and processed as indicated above.

### Mouse Xenograft Studies

All animal protocols related to mouse experiments were approved by the University of Chicago Institutional Animal Care and Use Committee. Approximately 1 x 10^6^ MDA-MB-231 cells in 100 μL of PBS were injected into the fourth mammary fat pad of 8–10-week-old female athymic nude mice (Charles River). Approximately 2.5 x 10^6^ U87-MG cells in 100 μL of 1:1 PBS/Matrigel (phenol red-free, Corning) were injected subcutaneously on the lower left flank of 16-week-old male athymic nude mice (Charles River). When tumors reached approximately 500mm^3^ in volume, mice were sacrificed, and tumors were dissected. The tumor samples were then gently diced, rinsed in DPBS, and incubated in RGDL (50 μM) in phenol-red-free RPMI media or vehicle for 1 hour with rotation at room temperature. Excess probe was removed, and the samples were then treated with BP (250 μM) in DPBS solution and irradiated at 440 nm for 1 or 5 min over ice. After irradiation, tissue samples were washed 4 times with 1 mL DPBS. Tissues were then treated with 1 x RIPA buffer with protease inhibitor and tip sonicated over ice. Lysates were then clarified and subjected to streptavidin enrichment and proteomic sample preparation as indicated.

### Whole proteome LC-MS/MS analysis of TNBC cells

MDA-MB-231 and BM1 cells were grown to confluence in 6 cm dishes before scraping and pelleting. Cells were washed with DPBS twice (1 mL) before resuspension in 4 M Urea and 50 mM ammonium bicarbonate with protease inhibitor tablet. Cells were then lysed via tip sonication over ice. Lysates (500 μl, each normalized to 1 mg/ml) were then treated with TCEP (10 mM final) and heated to 65 °C for 20 min. Samples were then cooled to room temperature before the addition of iodoacetamide (15 mM final) and incubation at room temperature for 30 min in the dark. Lysates were then diluted with a like volume of DPBS and CaCl_2_ (1 mM final) was added. Sequencing grade trypsin was then added (1:50 trypsin:protein) at 37 °C overnight. Peptide digest reactions were then stopped by cooling to room temperature and the addition of formic acid (2% final). Peptides were then desalted as indicated above. Peptides were then reconstituted in LC-MS grade water (0.1% Formic Acid, Optima) and normalized using Quantitative Colorimetric Peptide Assay (Pierce) before analysis by LC-MS/MS.

### LC-MS/MS Acquisition and Analysis

The proteomic methods reported are adopted from our previous reports^70^. LC-MS/MS analysis for proteomics samples was performed with an UltiMate 3000 RSLCnano System (Thermo Fisher Scientific) using an Acclaim PepMap RSLC C18 column (75 μm × 15 cm, 2 μm, 100 Å, Thermo Fisher Scientific) with an in-line Acclaim PepMap 100 C18 trap column (75 μm × 2 cm, 3 μm, 100 Å, Thermo Fisher Scientific) heated to 45 °C. The LC system was coupled to an Orbitrap Exploris 480 and Nanospray Flex Ion Source with stainless steel emitter tip (Thermo Fisher Scientific). Mobile phase A was composed of H_2_O supplemented with 0.1% formic acid, and mobile phase B was composed of CH_3_CN supplemented with 0.1% formic acid. The instrument was run at 0.3 μl min^−1^ with 2 h gradients. MS/MS spectra were collected for the entirety of the gradient using a data-dependent, 2-second cycle time setting with the following details: full MS scans were acquired at a resolution of 120,000, scan range of 380 *m*/*z* to 1,500 *m*/*z*, maximum IT of 25 ms, normalized AGC target of 300% and data collection in profile mode. MS2 scans were performed by high-energy collision dissociation (HCD) fragmentation with a resolution of 15,000, normalized AGC target of 50%, maximum IT of 50 ms, HCD collision energy of 30% and data collection in centroid mode. The isolation window for precursor ions was set to 1.6 *m*/*z*. Peptides with a charge state of 1, 7+ and unassigned were excluded, and dynamic exclusion was set to 40 seconds. The RF lens % was set to 40 with a spray voltage value of 2.0 kV and an ionization chamber temperature of 300 °C.

Data were processed using the SEQUEST HT search engine node within the Proteome Discoverer 3.0 software package. Data were searched using a concatenated target/decoy UniProt database of the human proteome with isoforms. Digest enzyme specificity was set to trypsin with up to two missed cleavages allowed, and peptide length was set to between 6 and 144 residues. Precursor mass range was set to 350–6500. Precursor mass tolerance was set to 10 ppm, and fragment mass tolerance was set to 0.02 Da. Up to 4 dynamic modifications were allowed per peptide, including heavy lysine (+8.0142), heavy arginine (+10.0083), oxidized methionine (+15.9949), N-terminal acetylation (+42.0106), N-terminal Met-loss (−131.0405) and N-terminal Met-loss + acetylation (−89.0299). Cysteine carboxyamidomethylation (+57.0215) was set as a static modification. A minimum of two peptides, with a minimum length of 6, was required for protein identification, and false discovery rate (FDR) was determined using Percolator with FDR rate set at 1%. Before quantification, chromatographic alignment was performed, with a maximum retention time difference of 10 min allowed, a mass tolerance of 10 ppm and a minimum signal/noise threshold of 5 required for feature mapping. SILAC ratios were determined using precursor-based quantification in a pairwise manner based on peak intensity without normalization or scaling using a maximum ratio of 20. For RGDL-PhotoPPI studies using SILAC for quantitation, proteins considered as enriched interactors were detected and quantified in at least 2 biological replicates and exhibited an RGDL-dependent median SILAC ratio greater than 4. MS files from LFQ samples were analyzed using the processing method parameters as indicated above with no dynamic modifications for heavy lysine or arginine. For whole proteome analysis of MDA-MB-231 and BM1 cells, a normalized pairwise ratio of BM1 over MDA-MB-231 was calculated in Proteome Discoverer. Proteins were considered significantly higher expressed if they displayed a fold change of at least 2 with a p-value less than 0.05. For RGDL experiments on tissue samples, abundance ratios of ‘probe’ over ‘no-probe’ were generated using Perseus accompanied by unpaired *t*-test. Positively enriched proteins were selected with the criteria of containing at least 4 peptides in the RGDL treated sample and a log_2_(ratio) greater than or equal to 4 with a *p*-value less than 0.05.

StringDB version 11.5 was used to find known PPIs among the enriched proteins^71^. Physical networks from PPIs with experimental evidence from literature and databases using medium confidence (interaction score of at least 0.400) were selected in StringDB searches. Scatter plots (referred to as “Yang plots” in the text) showing the relationship between known PPIs and grouped ion intensity of enriched proteins were generated from the node degrees data from STRING analyses (Y-axis values) and the median grouped ion intensity (X-axis values) across biological replicates within the experiment at hand. *Z*-scores for the enriched proteins were calculated using the median grouped ion intensity.

### Gene ontology and pathway analyses

Enriched proteins that are members of the GO term ‘Integrin Binding’ were used to generate integrin signaling sub-networks^72, 73^. Metascape analyses were performed on enriched proteins to identify significantly enriched gene sets and pathways^74^. Gene sets from GO Molecular Functions, GO Biological Processes, Hallmark Gene Sets and KEGG Pathway were used for Metascape queries^72, 75–77^. Enrichment P-values from the Metascape analyses were used to generate radar plots and heatmaps to compare differences and similarities between sample types. In addition, some significantly enriched pathways were selected for detailed heatmaps showing abundance ratios of proteins belonging to the respective pathway. Heatmaps were generated using Morpheus software (Broad Institute).

### Principal component analyses of abundance ratios of enriched proteins

Principal component analyses (PCA) were performed using the parallel analysis method within Graph Pad Prism 10 software. Briefly, protein abundance ratios from biological replicates were used as input data for PCA analysis. The data were centered to have a mean of 0, and principal components with eigenvalues greater than the 95^th^ percentile of the eigenvalues from 1000 simulations were selected. PCA loading plots showing PC1 and PC2 of every biological replicate were generated by Prism.

### Differential enrichment analyses of MDA-MB-231 and BM1 cells

Identified proteins from MDA-MB-231 and BM1 with positive log_2_(ratio) values were considered to compare differential enrichment between the two cell lines. A value of 0 was then assigned using imputation for missing log_2_(ratio) values. T-tests (two tailed, unpaired) on log_2_(ratios) of four biological replicates were performed in Perseus software to identify proteins with significantly different enrichment between the two cell lines (P-value < 0.01). Proteins with at least a 2-fold ratio difference were considered as higher enriched in one cell line over the other. For uniquely enriched proteins with a maximum ratio in each biological replicate, -log_10_(P-value) was uniformly set to 10. -Log_10_(P-value) from T-tests and log_2_(ratio_BM1_/ratio_231_) were used to generate a volcano plot.

Metascape analyses were then performed on these two groups: unique or higher enriched proteins in MDA-MB-231 cells (“231 high”) or BM1 cells (“BM1 high”).

The Proximity Score of a protein was calculated using the formula: Proximity Score = log_2_(abundance ratio) * log_10_(“Heavy” ion intensity). For each protein, a Proximity Score was calculated for every biological replicate using its respective abundance ratio and “heavy” ion intensity. Proteins with an abundance ratio less than 4 in both cell lines (i.e., not enriched in the RGDL experiment) were assigned a Proximity Score of 0.

### Survival analyses of breast cancer patients

Genes from proteins selected from the integrin interactome profiling data were used for patient survival analyses. A 20-gene signature was selected from MDA-MB-231 cells or BM1 cells based on the significant and unique enrichment of the target in either the 231 or BM1 RGDL-treated sample, high median ion intensity within that enriched target group and belonging to plasma membrane or cell surface gene ontology groups. For inclusion in the mRNA expression analysis the integrin-interacting target must be covered by the microarray probe set used in the existing patient survival dataset. Interaction barcode protein lists were identified as discussed in text and in Extended Data Fig. 4E; each gene list was used to query relapse-free survival data of TNBC patients from a microarray-based dataset or overall survival data of breast cancer patients without filtering of subtypes from an RNA-seq-based dataset^68^. TNBC patients were identified by all negative results in ER status by IHC, PR status by IHC, and ERBB2 status by microarray. Patient survival analyses were performed using the KM-plotter software^78^. Patients were grouped by average expression levels across each 20-gene signature, and the high and low-expression cutoffs were determined by KM-plotter by selecting the cutoff resulting in the highest statistical significance in the logrank test by *p*-value.

## Acknowledgements

We thank S. Ahmadiantehrani for her valuable suggestions during the writing process, figure drafting and proofreading assistance; M. Gardel, S. Seetharaman and M. Rosner for helpful discussions; M. Rosner and M. Henn for MDA-MB-231 tumor xenografts. This work was supported by NIH Multidisciplinary Training Grant in Cancer Research (MTCR) T32-CA09594 (to A.C.), The University of Chicago Department of Chemistry Josef Fried Chemical Biology Fellowship (to A.C), NIH Grant DP2GM128199-01 (to R.E.M.) and R01-GM145852 (to R.E.M), the Komen Career Catalyst Research CCR 21663985 and Alfred P. Sloan FG–2020–12839 (to R.E.M.).

## Author Contributions

A.C. and R.E.M. conceived of the study, probe design and the RGDL-PhotoPPI methods. A.C. and D.T. performed RGDL syntheses. A.C., S.H. and K.H. performed biochemical assays and cell-based assays. A.C. and D.Y. performed mass spectrometry and analysis of cellular and tissue RGDL-PhotoPPI experiments. A.C., D.Y. and R.E.M. performed data and bioinformatic analyses. R.E.M. supervised research. A.C, D.Y. and R.E.M wrote the manuscript with input from all authors.

## Additional Information

### Extended Data

Extended Data Figures 1-5 and Extended Data Tables 1-2.

### Supplementary Information

Synthetic materials, methods and characterization.

### Data Availability

Raw proteomics data files will be available in the MassIVE database upon publication.

### Correspondence or request for materials

should be addressed Raymond E. Moellering.

## Extended Data Figures

**Extended Data Fig. 1:**
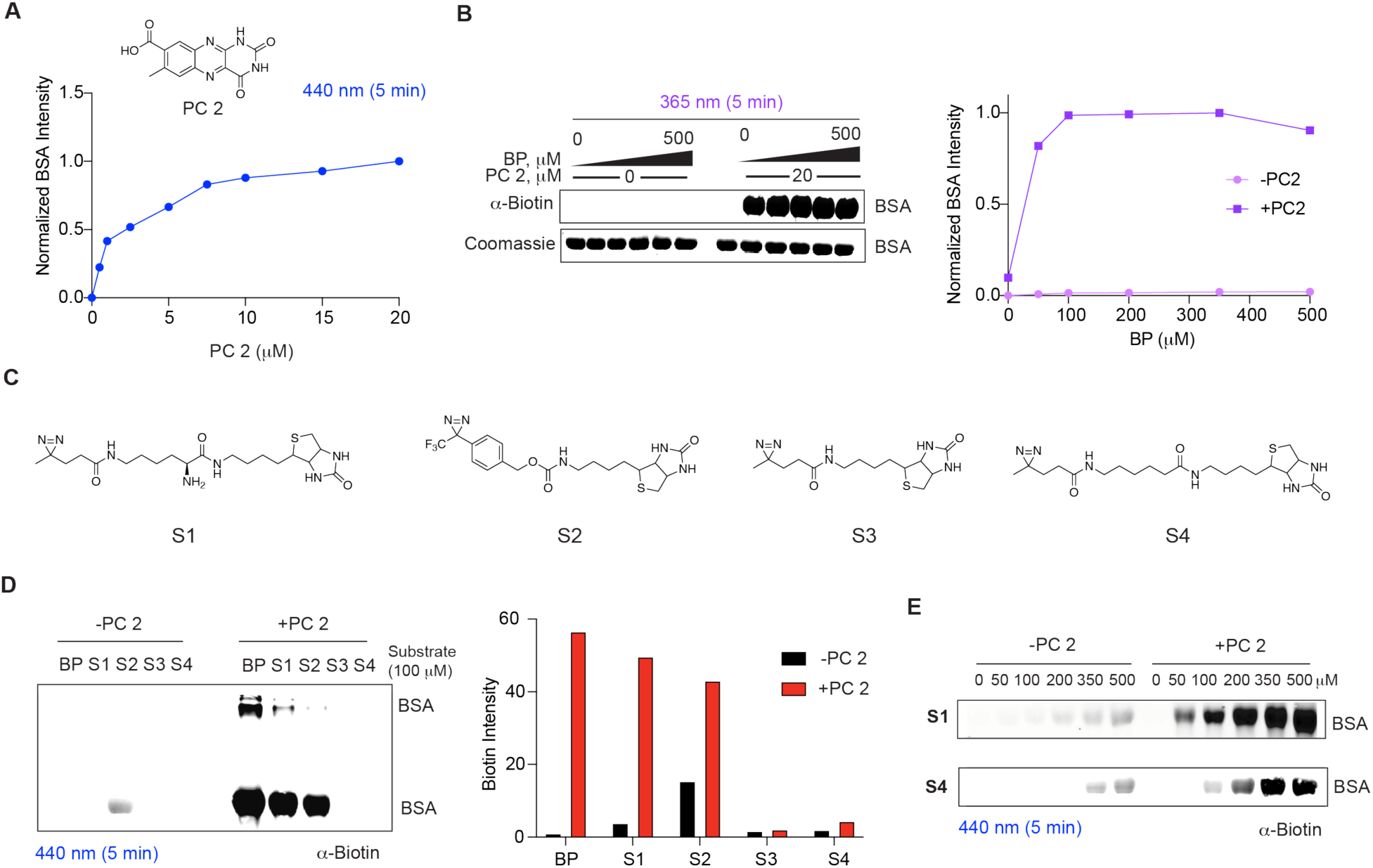
Lumichrome photosensitization of biotin-tagged substrates. **A**) Immunoblot demonstrating dose dependence of photocatalyst **2** (PC 2) on BP photosensitization. **B**) Immunoblot (*left*) and quantification (*right*) of BSA photolabeling with PC2 and BP at 365 nm. **C**) Chemical structures of substrate probes: S1, S2, S3 and S4. **D**) Screen of substrates S1-S4 photosensitization with PC 2; immunoblot (*left*) and quantification (*right*). **E**) Dose response of S1 (*top*) and S4 (*bottom*) in the presence of vehicle or PC 2, and 440 nm light.

**Extended Data Fig. 2:**
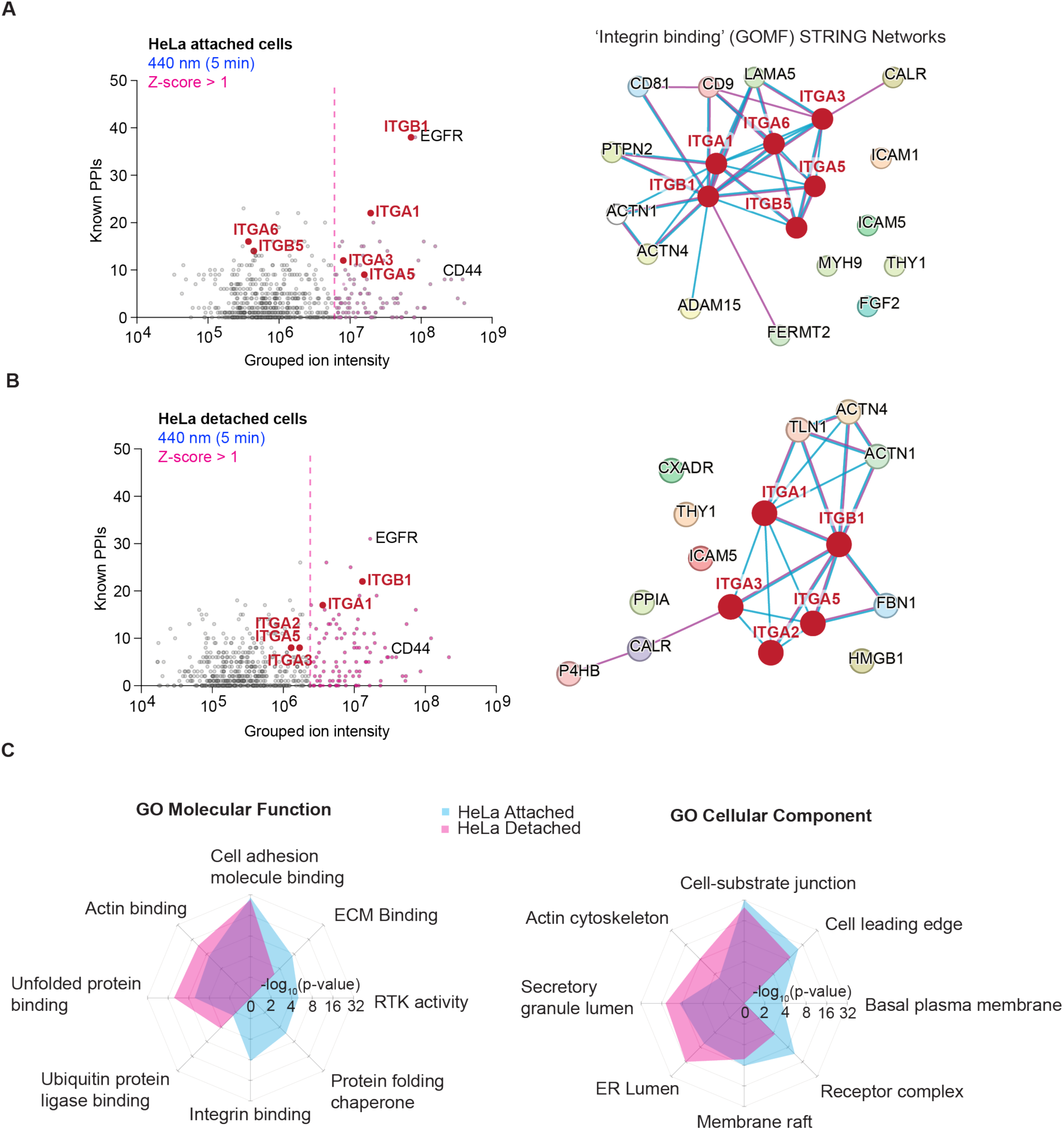
STRING analysis of HeLa cells in attached and detached cellular states. **A)** Yang plot of RGDL-enriched proteins (*left*) and GOMF ‘integrin binding’ STRING network of enriched proteins from attached HeLa cells (*right*); n = 4. **B)** Yang Plot of RGDL-enriched proteins (*left*) and GOMF ‘integrin binding’ STRING network of enriched proteins from detached HeLa cells (*right*); n = 3. **C)** Radial plots of GO Molecular function (*left*) and GO Cellular Component (*right*) terms from RGDL-PhotoPPI profiling of adhered and detached HeLa cells.

**Extended Data Fig. 3:**
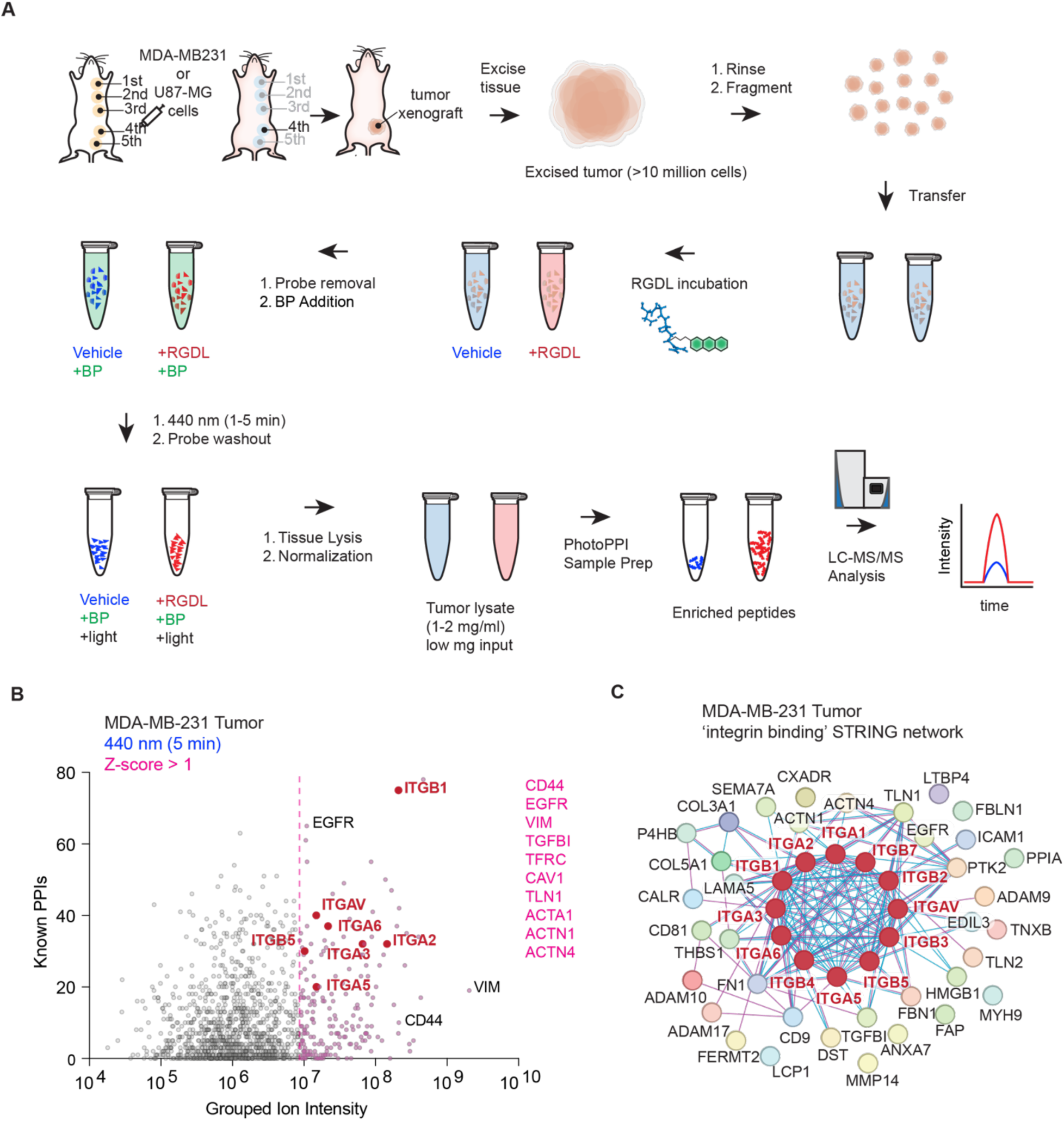
Rapid integrin interactome profiling of breast cancer-derived tumor xenografts. **A**) Schematic of RGDL-PhotoPPI tissue profiling workflow. **B**) Yang Plot of RGDL-enriched proteins from a MDA-MB-231 tumor xenograft; *n* = 1. **C**) GOMF ‘integrin binding’ STRING network of enriched proteins from MDA-MB-231 tumor xenograft.

**Extended Data Fig. 4:**
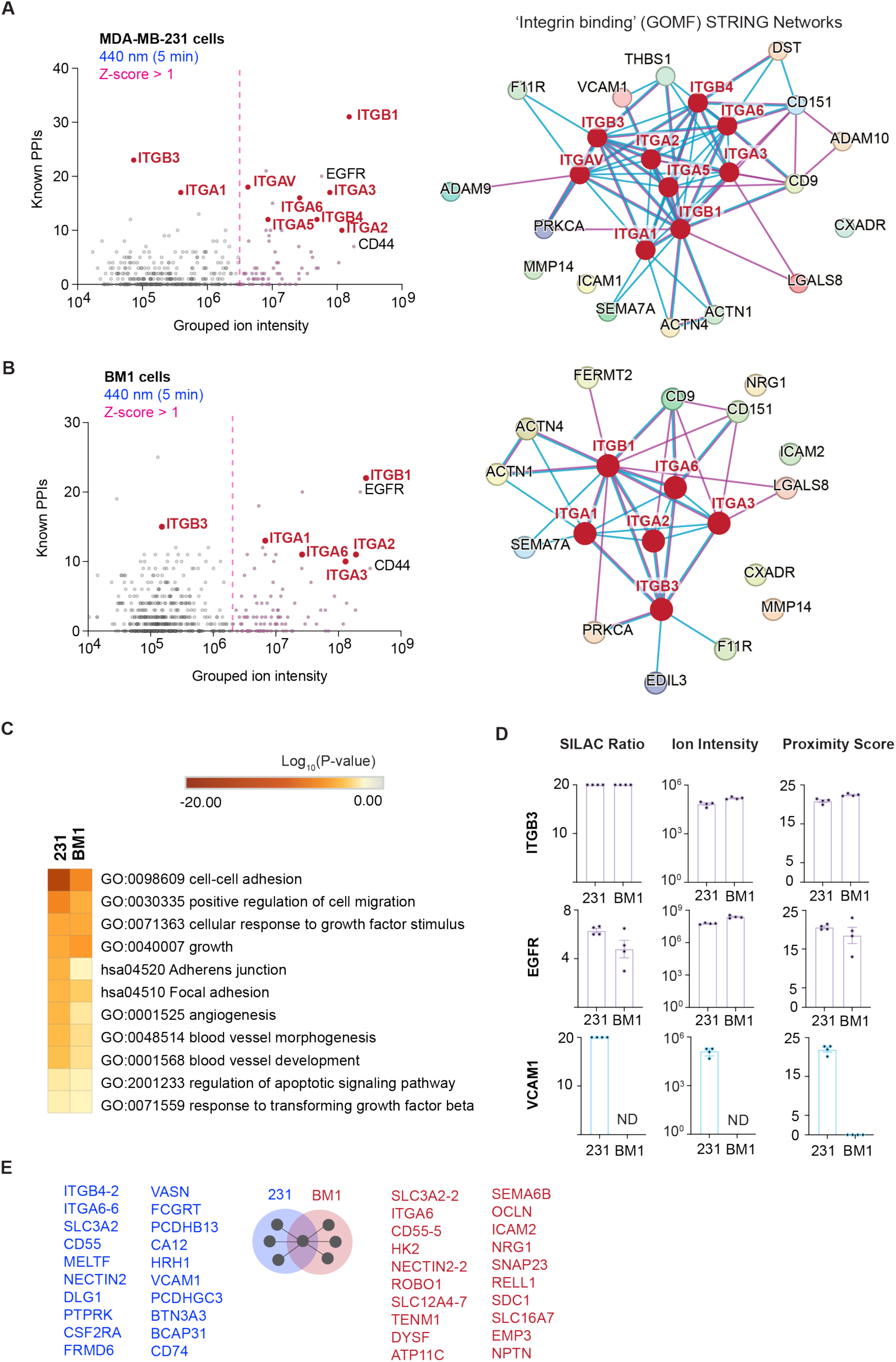
STRING analysis of RGDL-PhotoPPI profiling of MDA-MB-231 and BM1 cells. **A)** Yang Plot of RGDL-enriched proteins (*left*) and GOMF ‘integrin binding’ STRING network of enriched proteins from MDA-MB-231 cells (*right*); *n* = 4. **B)** Yang Plot of RGDL-enriched proteins (*left*) and GOMF ‘integrin binding’ STRING network of enriched proteins from BM1 cells (*right*); *n* = 4. **C)** Heatmap of unique GO, KEGG, and HALLMARK pathway terms from Metascape analysis of the data in (A-B). **D**) Bar plots of SILAC ratio, median RGDL ion intensity and proximity scores for ITGB3, EGFR and VCAM1. Data represents mean ± S.E.M. from *n* = 4 biological replicates. **E**) 20 gene survival signature lists for MDA-MB-231 (blue) and BM1 (red).

**Extended Data Fig. 5:**
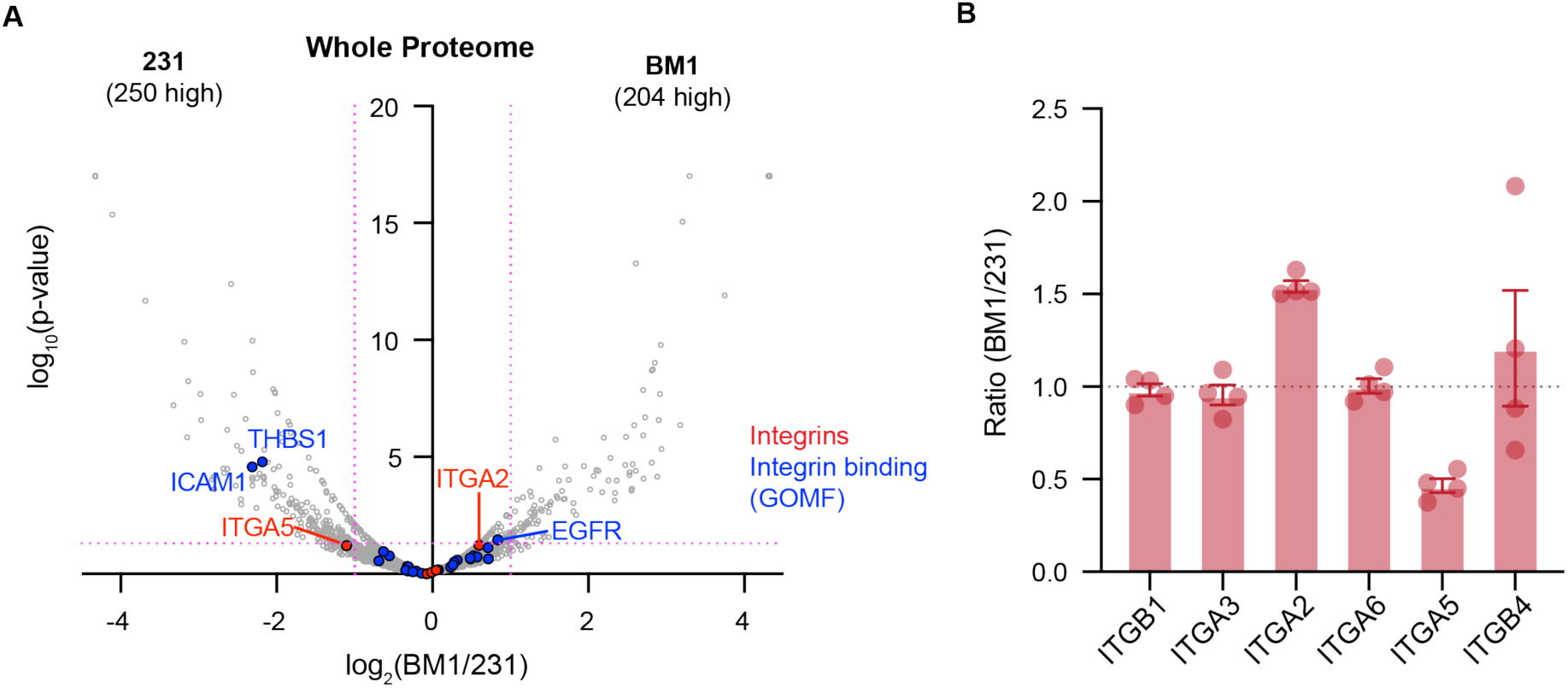
Whole proteome analysis of MDA-MB-231 and BM1 cells. **A**) Volcano plot of proteome abundance comparison of MDA-MB-231 and BM1 cells from n = 4 biological replicates. Integrins are highlighted in Red and ‘integrin binding’ (GOMF) proteins are highlighted in Blue. Cutoffs for ratio and p-value in Magenta indicate 2-fold change and p-value of 0.05. **B**) Plot of ratios for integrins detected and quantified in A.

